# Persistent Cyfip1 expression is required to maintain the adult subventricular zone neurogenic niche

**DOI:** 10.1101/781856

**Authors:** Christa Whelan Habela, Ki-Jun Yoon, Namshik Kim, Arens Taga, Kassidy Bell, Dwight E. Bergles, Nicholas J. Maragakis, Guo-li Ming, Hongjun Song

**Author notes:** Corresponding Author: Hongjun Song, Ph.D., Perelman School of Medicine at University of Pennsylvania, Department of Neuroscience, 415 Curie Blvd, Suite 111B, Philadelphia, PA 19104, USA. Co-Corresponding Author: Christa Whelan Habela, M.D, Ph.D., Johns Hopkins Medicine, Department of Neurology, 600 North Wolfe Street, Meyer 2-147, Baltimore, MD 21287, USA. Author contributions: C.W.H. led the project, collected data and was involved in all aspects of the study. K-J.Y. and N.K. generated the Cyfip1 floxed animals and planned breeding paradigms. K.B. assisted with immunostaining and animal breeding. A.T. contributed to data analysis. G-l.M., N.J.M. and D.E.B contributed to analysis, experimental design and writing. C.W.H., KJ-Y, G-l.M. and H.S. conceived of the project and experimental design. G-l.M. and H.S. oversaw the project. C.W.H. and H.S. wrote the manuscript.

## Abstract

Neural stem cells (NSCs) persist throughout life in the subventricular zone (SVZ) niche of the lateral ventricles as B1 cells. Maintaining this population of NSCs depends on the balance between quiescence and self-renewing or self-depleting proliferation. Interactions between B1 cells and the surrounding niche are important in regulating this balance, but the mechanisms governing these processes have not been fully elucidated in adult mammals. The cytoplasmic FMRP-interacting protein (CYFIP1) regulates apical-basal polarity in the embryonic brain. Loss of Cyfip1 during embryonic development in mice disrupts the embryonic niche and affects cortical neurogenesis. However, a direct role for Cyfip1 in the regulation of adult NSCs has not been established. Here, we demonstrate that Cyfip1 expression is preferentially localized to B1 cells in the adult SVZ. Loss of Cyfip1 in the embryonic mouse brain results in altered adult SVZ architecture and expansion of the adult B1 cell population at the ventricular surface. Furthermore, acute deletion of *Cyfip1* in adult NSCs results in a rapid change in adherens junction proteins as well as increased proliferation and the number of B1 cells at the ventricular surface. Together, these data indicate that CYFIP1 plays a critical role in the formation and maintenance of the adult SVZ niche and, furthermore, deletion of Cyfip1 unleashes the capacity of adult B1 cells for symmetric renewal to increase the adult NSC pool.

**SIGNIFICANCE:** Neural stem cells (NSCs) persist in the subventricular zone (SVZ) of the lateral ventricles in adult mammals and their population is determined by the balance between quiescence and self-depleting or renewing cell division. The mechanisms regulating their biology are not fully understood. This study establishes that the cytoplasmic FMRP interacting protein 1 (Cyfip1) regulates NSC fate decisions in the adult SVZ and NSCs that are quiescent or typically undergo self-depleting divisions retain the ability to self-renew in the adult. This contributes to our understanding of how adult NSCs are regulated throughout life and has potential implications for human brain disorders.

## INTRODUCTION

Neural stem cells (NSCs) persist in the subventricular zone (SVZ) of the lateral ventricles into adulthood in mammals (Doetsch et al., 1999a; Bond et al., 2015; Altman, 1969). The adult SVZ recapitulates the developmental NSC niche with an apical – basal polarity of the NSCs, referred to as type B1 cells (Lois and Alvarez-Buylla, 1993; Doetsch et al., 1999b). The cell bodies of B1 cells lie beneath the ependymal layer and undergo symmetric self-renewing divisions to maintain their population or self-depleting divisions to generate olfactory bulb interneurons and oligodendrocyte precursors (Parras et al., 2004; Menn et al., 2006; Rousselot et al., 1995; Lois and Alvarez-Buylla, 1994; Lois and Alvarez-Buylla, 1993; Obernier et al., 2018). Disruption of the SVZ niche leads to altered proliferation as well as altered neuronal and oligodendrocyte genesis (Kokovay et al., 2012; Jimenez et al., 2009; Relucio et al., 2012). The structure of the niche changes with age as B1 cells are depleted (Shook et al., 2012; Obernier et al., 2018), but the cellular mechanisms regulating niche maintenance and B1 cell fate in the adult brain are still not fully elucidated.

Type B1 cells project apical processes to the ventricular surface and basal processes to the vasculature underlying the adult SVZ. At the ventricle, the apical processes are surrounded by ependymal cells forming an epithelial surface and oriented in a pinwheel-type formation around the apical processes (Mirzadeh et al., 2008; Doetsch et al., 1999b; Alvarez-Buylla and Lim, 2004; Mercier et al., 2002). Central to the niche structure in both the embryo and the adult is the maintenance of apical-basal polarity (Lian and Sheen, 2015a; Bizzotto and Francis, 2015; O’Leary et al., 2017; Yoon et al., 2014). During embryonic development, radial glial cells make apical connections to the ventricular surface and basal projections to the pia (Gotz and Huttner, 2005; Kosodo et al., 2004) and the orientation of the division plane along the apical-basal axis regulates the fate of the daughter cells (Kosodo et al., 2004). This polarity is determined by the presence of adherens junctions and loss of junction integrity in fetal development leads to alterations in cellular polarity and abnormal neural development (O’Leary et al., 2017; Bizzotto and Francis, 2015; Ferland et al., 2009; Guerra et al., 2015; Lian and Sheen, 2015b; Yoon et al., 2014).

The stability of adherens junctions is dependent on cadherins interacting with the cytoplasmic actin ring (O’Leary et al., 2017; Priya and Yap, 2015; Verma et al., 2012). This is mediated by Arp2/3 dependent actin nucleation and the WAVE regulatory complex (WRC) (Verma et al., 2012; Wang et al., 2016). Cytoplasmic FMRP interacting protein1 (Cyfip1) interacts with Rac-GTP to cleave the WRC, resulting in actin polymerization. Cyfip1 regulates apical-basal polarity and loss of the protein in embryonic development results in adherens junction deficits (Yoon et al., 2014). Adult mice that are haploinsufficient for Cyfip1 have impaired myelination and decreased oligodendrocytes in the corpus callosum as well as behavioral abnormalities (Silva, A. I., Haddon et al., 2019; Dominguez-Iturza et al., 2019).

In this study we show that there is persistent expression of Cyfip1 in type B1 cells of the adult SVZ in mice with prominent localization to the apical processes projecting to the ventricular surface. Deletion of *Cyfip1* during embryonic development results in an expansion of the B1 cell population, altered localization and increased proliferation rates in the adult SVZ. Acute loss of *Cyfip1* in the adult SVZ NSCs is sufficient to alter the localization and increase proliferation rates of the B1 cells, suggesting that Cyfip1 suppresses symmetric B cell expansion in the adult. Changes in adherens junction protein localization parallels decreases in Cyfip1 expression and supports the hypothesis that there is an underlying loss of adherens junction stabilization.

## MATERIALS AND METHODS

### Animals

All transgenic animals were crossed on a C57/Bl6 background. The *Nestin-CreER* animals were kindly provided by Gordon Fishell (Balordi and Fishell, 2007). *Nestin-Cre* (JAX stock #003771 – B6.Cg-Tg(Nes-cre)1Kln/J) (Giusti et al., 2014; Tronche et al., 1999) and *mTmG* reporter mice (Stock # 007676 B6.129(Cg)-Gt(ROSA)26Sortm4(ACTB-tdTomato,-EGFP)Luo/J) (Muzumdar et al., 2007) were obtained from the Jackson Laboratory (Bar Harbor, ME).

In order to generate a Cyfip1 floxed allele (*cyfip1*^*f*^), a targeting vector was designed to insert a loxP sequence in front of exon 2 as well as a positive selection marker (PGK promoter driven neomycin resistant gene) together with another loxP sequence next to exon 5. This was constructed by recombineering as described (Liu et al., 2003). Specifically, an 11.9 kb genome fragment containing exon 2 to exon 5 from 129Sv BAC clone (bMQ182K14, Source Bioscience) was retrieved into a PL253 plasmid containing a negative selection marker (MC1 promoter driven thymidine kinase gene) using homologous recombination. A loxP sequence and an Flpe-PGK-EM7-Neo-Flpe-loxP cassette were sequentially inserted into the engineered PL253, resulting in 6.0 kb and 1.0 kb homology arms. The targeting vector was linearized and electroporated into 129S4/Sv Jae embryonic stem cells (The Transgenic Core Laboratory in Johns Hopkins School of Medicine), and homologous recombination was confirmed by PCR screening. Targeted clones were injected into C57BL/6J blastocysts, which were subsequently transferred into pseudo-pregnant foster mothers. Confirmation of germ-line transmission of the floxed allele and routine genotyping were performed by PCR screening on tail genomic DNA (wt, 470bp; floxed, 520bp) using DNA primers as follows: 5’-GCACCTCTCTGCATTTCTGT-3’ and 5’-GCACCAATCAAGTGTTTTCC-3’.

For conditional knockout experiments, homozygous *Cyfip1*^*f/f*^ animals were crossed and back crossed with animals heterozygous for *Nestin-Cre to generate Nestin-Cre*:*Cyfp1*^*f/f*^ males that were heterozygous for *Nestin-Cre* with homozygous floxed *Cyfip1* alleles. These were subsequently bred with *Cyfp1*^*f/f*^ females resulting in 50% control (*Cyfp1*^*f/f*^) and 50% conditional knockout (cKO) animals (*Nestin-Cre:Cyfp1*^*f/f*^). Inducible breeding pairs were made up of *Nestin-CreER:Cyfip1*^*+/f*^:*mTmG* males bread with *Cyfip1*^*+/f*^:*mTmG* females. The mTmG allele was either heterozygous or homozygous in the experimental animals.

All experiments involving animals were approved by the animal care and use committee at Johns Hopkins University. Both male and female animals were used for experiments. Animals were housed under 14 hour light / 10 hour dark housing conditions with standard diets and water *ad libitum*.

### Immunocytochemistry

Anaesthetized animals were perfused with phosphate buffered saline (PBS) followed by 4% paraformaldehyde (PFA). Brains were removed from skulls and placed in 4% PFA overnight and no longer than 24 hours at 4°C. They were then washed one time with PBS and placed in a 30% sucrose in PBS solution at 4°C for a minimum of 48 hours prior to cutting. Serial coronal brain sections were collected using a sliding microtome (Leica, SM2010R) or a cryostat (ThermoFisherHM 505 and an HM 550) after brains were frozen in either 30% sucrose solution or OCT freeze solution (Sigma). Sections were stored frozen in multi well plates containing antifreeze solution (300 g sucrose, 300 ml ethylene glycol, 500 ml 0.1 M PBS). Prior to antibody staining, anti-freeze solution was removed and sections were washed 2 times with PBS. Antibody solutions were made up of 5% donkey or goat serum, 3% bovine serum albumin (BSA) and 0.05% Triton X 100 in PBS or TBS. Primary antibodies were incubated at 4°C for 24 to 72 hours. Sections were washed 3 times in 0.05% Triton-X 100 in PBS solution prior to secondary antibody application. Secondary antibodies were diluted in the above-described antibody solution using Alexa Fluor 488, 555, 568, and 647 secondary antibodies (ThermoFisher Scientific) at 1:400 dilutions or donkey serum for donkey Cy2, CY3 and Cy5 antibodies (Jackson ImmunoResearch) at 1:250 dilution. Secondary antibody solutions were incubated either at room temperature for 2 to 4 hours or overnight at 4°C. Hoechst 33342 (Sigma) or DAPI (Roche) nuclear stains were added to the secondary antibody solutions. Brain sections that were stained with antibodies that required antigen retrieval were incubated in Dako 1X target retrieval solution (Agilent Dako) or sodium acetate buffer, pH 6 (Sigma) at 95 °C for 20 minutes then room temperature for 20 minutes prior to staining. If green fluorescent protein (GFP) staining was required, anti-GFP primary and secondary antibody staining was conducted prior to the antigen retrieval step. Tissue was mounted on Superfrost™ or Superfrost Plus™ slides (Fisher) and coverslipped with 2.5% PVA/DABCO mounting media (Sigma) or ProLong Antifade mounting media (ThermoFisher). Specific antibodies are noted in the results section and include: mouse anti β-Catenin (BD Biosciences Cat # 610153), mouse anti-ɣ-Tubulin (Abcam cat # ab11316), rabbit anti GFAP (Dako, Cat # Z0334), rabbit anti-Cytoplasmic interacting protein 1 (Millipore-Sigma, Cat # Ab6046), rabbit anti-β-Catenin polyclonal antibody (ThermoFisher, PA5-16762), chicken anti-Green Fluorescent Protein (Aves, Cat # NC9510598), mouse anti-N-Cadherin (Invitrogen, 981235A), rabbit anti – S100β (Sigma, HPA015768), mouse anti – S100β (Sigma, AMAB91038), goat anti – Sox2 (Santa Cruz Biotechnology, SC17320).

### Whole Mount Preparation

Whole mount preparations of the ventricular wall were obtained using a protocol modified from that published in extensive detail by Mirzedah et al (Mirzadeh et al., 2010). The one modification made for our experiments was that the animals were perfused with 4% PFA prior to starting the dissection rather than afterwards. Immunostaining of the whole mount sections was performed as described above.

### Cell Proliferation Quantification

Cells undergoing DNA replication in S phase were marked by incorporation of 5-ethynyl-2’-deoxyuridine (EDU) (Sigma, St. Louis, MO. Cat # 900584-500MG). A stock concentration of 32.5 mM EDU was made by adding EDU to sterile saline solution with the addition 1:1000 5 M NaOH and heating to 42°C for 30 to 60 minutes to dissolve. Stock solutions were stored at −80°C. Two to twenty four hours prior to perfusion, the solution was warmed to 37°C and intraperitoneal injections were conducted on 56-84 day old animals for a final dose of 200 mg / kg. After perfusion and antibody staining, the standard commercial protocol for the Click-iT™ Plus EDU cell proliferation kit for Imaging (ThermoFisher Scientific, Waltham, MA. cat. No. C10639) was used to fluorescently label the EDU incorporated into the newly synthesized DNA. 3-dimensional (3D) tiled images were then obtained of the cells and images were reconstructed either in Imaris 3D software (Bitplane), Image J Software or ZEN software (Carl Zeiss Microscopy, Jena, Germany). Cells were manually quantified based on the presence of EDU florescence in the cell nuclei.

A monoclonal antibody (Mouse-anti-MCM2, BD Biosciences, Cat#610701; RRID: AB_398024) directed against the minichromosome maintenance complex component 2 (MCM2) protein, that is involved in the onset of DNA replication and cell division (Mincheva et al., 1994) was used to mark cells within the cell cycle in animals that were not injected with EDU. Antibody labeling was detected by immunofluorescence after secondary antibody labeling. MCM2 expressing cells were quantified in the same manner as EDU.

### Tamoxifen Injection

A stock solution of 66.7 mg/mL of tamoxifen in a 5:1 solution of corn oil and ethanol was prepared. In order to dissolve the tamoxifen in the corn oil and ethanol solution, it was heated to 37°C with intermittent vortexing. Stock concentrations were stored at −80°C. Prior to use, tamoxifen was warmed to 37°C and then injected into the intraperitoneal space of P56 to P84 *Nes-CreER:mTmG* animals with or without the floxed *Cyfip1* trans-genes at a final concentration of 248 mg/kg. Animals underwent intracardiac perfusion 2 to 8 days post injection.

### Image Acquisition, Processing, and Quantification

Brain sections were imaged on either a Zeiss LSM 800, a Zeiss LSM 710, or a Zeiss 800 Airyscan confocal microscope (ZEISS International) on Zen Software (ZEISS International). Low magnification images were acquired with 10X or 20X air objectives. High magnification images were acquired with 40X or 63X oil immersion objectives. Z-stacks were obtained using the optimal inter-slice distance for the objective. For quantitative and qualitative experiments in which a control and an experimental condition were being compared, equal settings of laser intensity, pinhole aperture, and inter slice distance for Z-stacks were maintained as constant between conditions within the same experiment whenever possible. For larger fields of view multiple tiled sections were obtained and stitched together prior to exporting for analysis. 3D reconstructions were generating using Imaris Software 7.6 (Bitplane). Quantification of fluorescence intensity was measured in Adobe Photoshop (Adobe) or Image J software (NIH). Quantification of the number of cells expressing cell markers was determined using Imaris 7.6, Zen, or Image J software. Image preparation for publication was conducted in Adobe Photoshop (Adobe). Any modifications to brightness or contrast of images was applied equally to control and experimental images.

### Quantification and Statistical Analysis

All data are presented as the mean ± SEM. The n values refer to the number of animals used. In cases where multiple coronal sections were analyzed per animal, the average of these sections was determined for each n in the experimental population. The number of sections per animal used is indicated in the text and figure legends and a minimum of 3 sections per animal was used. Quantification was performed by a person who was blinded to the animal genotype at the time of quantification for all figures. Image acquisition was performed by a blinded person for Figures 2 and 3. Statistical analysis was performed using either Origin Pro 8.5 (OriginLab) or GraphPad Prism 7 (GraphPad Software Inc.). For statistical analysis experiments with only 2 conditions, a 2 way Students T test was used. Unless otherwise noted, data was unpaired. For comparisons between most multiple groups, a one way Anova followed by a T test was used. Sample sizes were not predetermined using statistical methods. P values reported were * for p < 0.05, ** for p < 0.01 and *** for p < 0.001.

**Figure 1.**
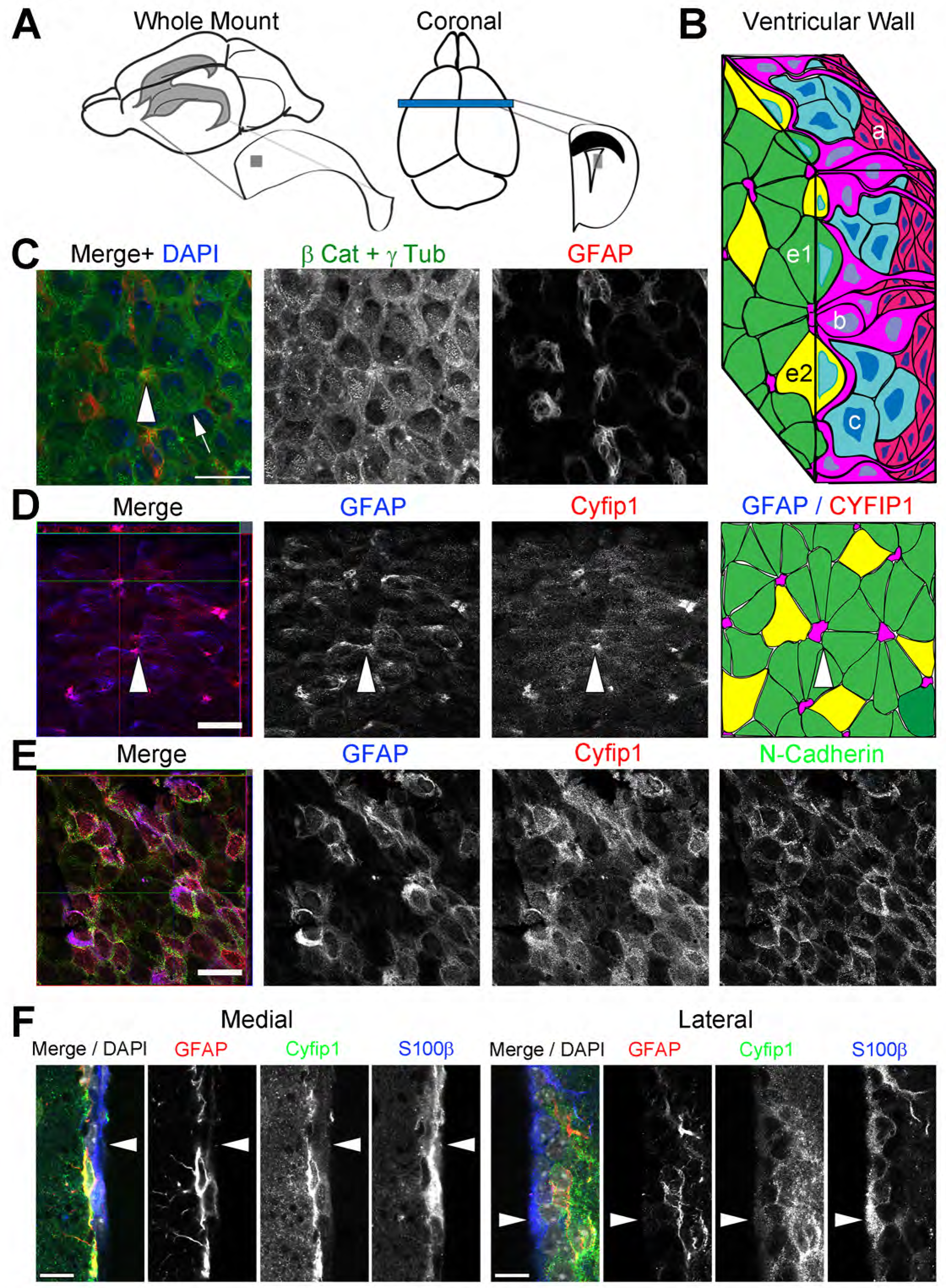
Cyfip1 is expressed in the B1 cells of the adult subventricular zone. (A) Diagrammatic description of the whole mount and coronal preparations used for analysis in this study. The gray squares correspond to the three-dimensional image in (B). (B) Three-dimensional diagram of the cellular composition of the adult subventricular zone. e1 = E1 ependymal cells. e2 = E2 ependymal cells. b = type B cells. c = type C transient amplifying cells. a = migratory neuroblasts. Model based on that of Mirzadeh et al. 2008. (C) Whole mount preparation of control ventricular surface with immunofluorescent staining for β-catenin and ɣ-tubulin (green) and GFAP (red). (D) Immunofluorescent staining of GFAP (blue) and Cyfip1 (red) at the ventricular surface with pictorial representation in the last panel. Arrowheads indicate apical projections at the center of the pinwheel. (E) GFAP and Cyfip1 expression 5 µm below the surface of the ventricular zone with N-cadherin ummunostaining in green. (F) Coronal sections with immunofluorescent staining for GFAP (red), Cyfip1 (green), S100 β (blue) and DAPI (grey). Images are examples from the medial and the lateral ventricular wall. Arrowheads indicate S100β^+^ Cyfip1^−^ ependymal cells. All images acquired with a 63X objective. Scale bars, 20 µm (C-E) and 10 µm (F). All images are representative of similar immunostaining seen in a minimum of 4 animals.

**Figure 2.**
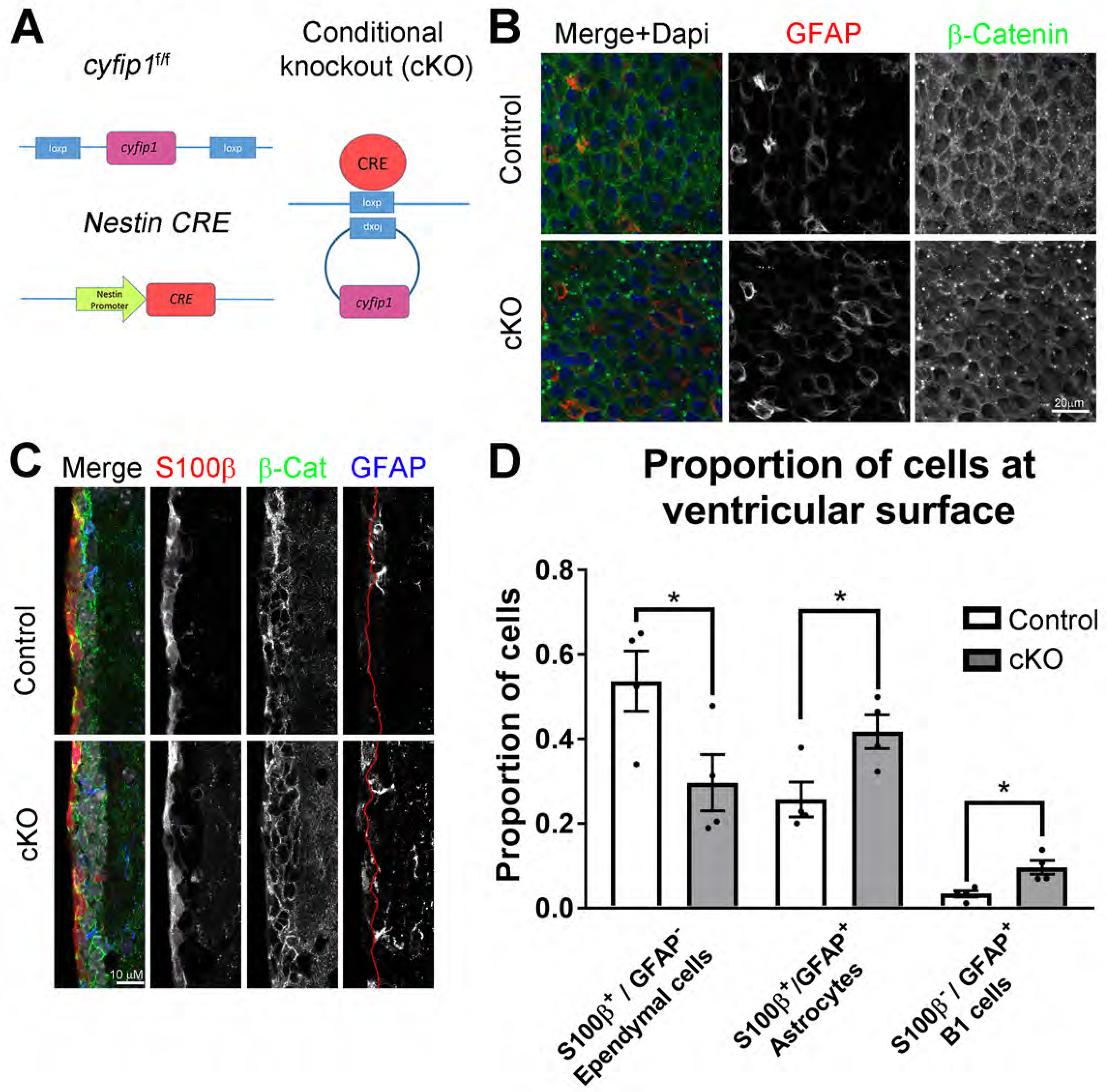
Loss of Cyfip1 alters the structure of the ventricular surface in adult Cyfip1 conditional knockout mice. (A) Schematic illustration of *the Nestin-Cre:Cyfip1*^*f/f*^ conditional knockout (cKO) animals. (B) Example images of the ventricular surface in adult mice. Whole mount preparations were immunostained for β-catenin (green) and GFAP (red). Images obtained with a 63X objective. Scale bar, 20 µm. (C) Example coronal images of the lateral ventricle of the adult SVZ in control versus cKO animals. Sections immunostained with antibodies targeting S100β (red), β-catenin (green), and GFAP (blue). Red line demarcates the border between the first cell layer at the ventricular surface and the SVZ zone. Images obtained with a 63X objective. Scale bar, 10 µm. (D) Quantification of the cellular make-up at the ventricular surface. The number ofGFAP and S100β expressing cells were quantified in relation to the total number of cells at the ventricular surface based on nuclear DAPI staining. Values represent mean ± s.e.m. (n = 4 control and 3 cKO animals. 3 sections per animal; * p < 0.05; Student’s t-test).

**Figure 3.**
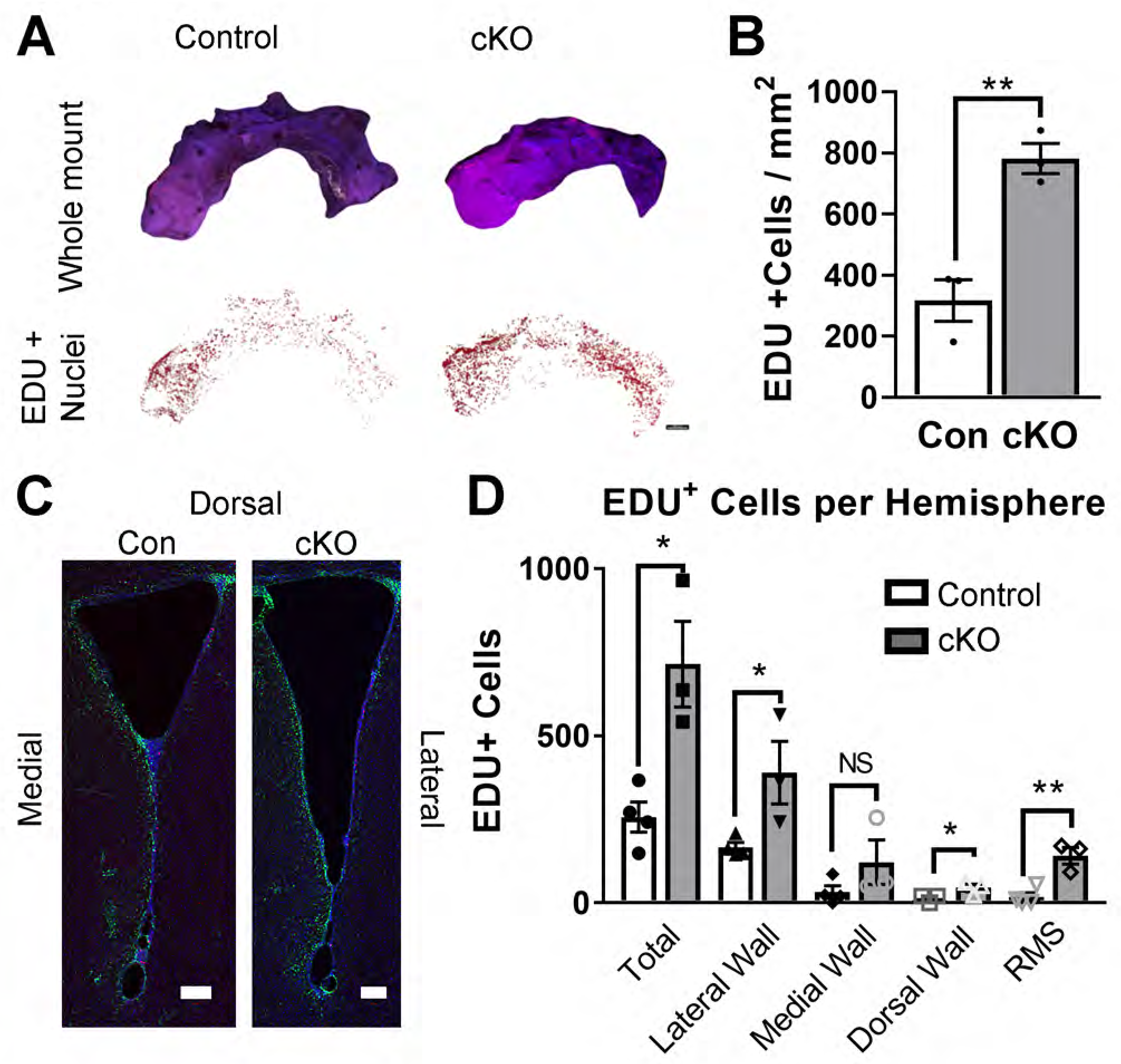
Loss of Cyfip1 in the adult subventricular zone results in altered proliferation. (A) Whole mount preparations of EDU-injected control and conditional knockout (cKO) mice 24 hours post injection. EDU was labeled using a Click-it reaction and tiled three-dimensional images were obtained using a fluorescent 10 X objective to capture the entire ventricular surface. Images were reconstructed in *Imaris* software and EDU^+^ nuclei were marked. Scale bar, 500 µm. (B) EDU^+^ cells as in (A) were quantified and normalized to total area of the subventricular zone for each animal (n = 3 animals for each condition). (C) Example images of coronal sections from control and cKO animals stained with EDU (red), GFAP (green), and DAPI (blue). Three-dimensional tiled images were obtained using a 20 X objective. Scale bars, 100 µm. Note different size of each scale bar. (D) Quantification of EDU^+^ cells from coronal sections. We examined 40 µm sections beginning from the posterior frontal lobe just anterior to the ventricle (rostral migratory stream, RMS) to the dentate gyrus of the hippocampus and examined every 6^th^ coronal section. For each animal, a minimum of 7 sections along the anterior to posterior axis of the lateral ventricles were examined. Values represent mean ± s.e.m. (n = 4 animals for control and 3 animals for cKO; **p < 0.01; *p < 0.05; Student’s t-test).

## RESULTS

### Cyfip1 expression persists in the neurogenic niche of the adult subventricular zone

To determine whether Cyfip1 is persistently expressed in the neurogenic niche of the adult SVZ zone, we examined whole mount preparations as well as coronal sections from C57/Bl6 mice between postnatal day 56 and 70 (Figure 1A). The SVZ niche at this age is characterized by a unique organization with central apical projections of type B1 cells to the ventricular surface forming the hub of the neurogenic niche architecture (Figure 1B) (Mirzadeh et al., 2008). When viewed *en face* from the ventricular surface in whole mount preparations (Figure 1B-D, surface; and 1E, 5 µm below the surface), the cells of the SVZ are organized into a “pinwheel” structure. The type B1 cells express the intermediate filament protein glial fibrillary acidic protein (GFAP) (Garcia et al., 2004). The apical processes of the GFAP^+^ B1 cells (Figure 1B, “b” and Figure 1C, arrowhead) are surrounded by epithelial-like ependymal cells with cilia comprised of γ-tubulin (Figure 1B, e1 and e2, Figure 1C, arrow). Cell-cell junctions are demarcated by β-catenin or N-cadherin localized to the adherens junctions (Figure 1C and 1E). The cell bodies of the B1 cells lie beneath the surface of the ventricular zone in the SVZ (Figure 1B and 1E). Immunostaining for Cyfip1 demonstrates that Cyfip1 is persistently expressed in the SVZ of the adult mouse (Figure 1D-F). It is expressed at the highest levels in the B1 cells and is localized to the apical processes of the B1 cells at the surface (Figure 1D, arrowheads) as well as the cell bodies of B1 cells below the surface (Figure 1E). It is localized in GFAP expressing cells in discrete clusters at the surface (Figure 1D, arrowheads). Below the ventricular surface, Cyfip1 staining is present in the cell bodies of GFAP^+^ cells and overlaps with N-cadherin staining at cell membranes (Figure 1E). In contrast, there is almost no Cyfip1 expression in the S100β^+^ GFAP^−^ ependymal cells at the ventricular surface (Figure 1F, arrowheads) and very low expression in the double positive S100β^+^ GFAP^+^ cells at the surface (Figure 1F), which likely represent mature astrocytes. This indicates that not only is Cyfip1 persistently expressed, it is specifically restricted to the GFAP^+^ S100β^−^ B1 cells of the adult SVZ.

### Cyfip1 is required for normal SVZ architecture and loss of Cyfip1 expression alters the cellular composition of the ventricular surface in adult

Persistent expression of the Cyfip1 protein in the adult SVZ and the preferential localization of Cyfip1 to the B1 type cells at the center of the pinwheel niche suggests that it continues to be important for the regulation of B1 cells in the adult niche. Germline deletion of C*yfip1* is embryonic lethal (Pathania et al., 2014). Therefore, we generated a conditional knockout animal using a *Cre-lox* system in which Cre expression is driven by the *Nestin* promoter, which is active in NSCs and progenitor cells of the neurogenic zones that persist into adulthood, including the SVZ (Giusti et al., 2014). In order to specifically delete *Cyfip1* in Nestin expressing neuronal precursors and their progeny, *Nestin-Cre* animals were crossed with *Cyfip1*^f/f^ animals resulting in homozygous deletion of the gene in Nestin expressing cells (Figure 2A).

Examination of the lateral ventricular surface of the *Nestin-cre*:*Cyfip1*^f/f^ conditional knockout animals (cKO) compared to littermate controls (Con) carrying the *Cyfip1*^f/f^ alleles, but not expressing Cre, reveals significant changes in the organization of the cells at the ventricular surface (Figure 2B). Namely, in whole mount sections, there appears to be an increase in the number of GFAP^+^ cell bodies at the ventricular surface of the cKO animals compared to the controls. Additionally, compared to the control SVZ, where there is prominent GFAP immunoreactivity at the apical processes of neural stem cells, and ventricular-projecting processes are not as clearly demarcated. There is also a change in β-catenin expression at the ventricular surface with less uniform staining at cell-cell junctions (Figure 2B).

We examined coronal sections of control and cKO animals in order to more clearly define the changes in cellular composition and organization seen in the whole mount preparations. Immunostaining with antibodies directed towards GFAP, S100β, and β-catenin demonstrates that there is an increase in the GFAP^+^ cell bodies of the NSCs at the ventricular surface (Figure 2C). We quantified the number of S100β and GFAP expressing cells at the cell surface as well as the number of cells that immunostained positive for both. In the control animals, the majority of the cells along the ventricular surface are S100β^+^ GFAP^−^ ependymal cells (Figure 2C and 2D). There is a significant decrease in the number of S100β^+^ GFAP^−^ cells relative to the total cells at the ventricular surface based on DAPI staining in the cKO animals compared to the control aniamls (Con = 0.53 ± 0.07 S100β^+^ GFAP^−^ cells / total cells, n = 4 vs cKO = 0.30 ± 0.07 S100β^+^ GFAP^−^ cells / total cells, n = 4; p < 0.05, unpaired T-Test) that is paralleled by an increase in the number of S100β^+^ GFAP^+^ mature astrocytes (Con = 0.26 ± 0.04 S100β^+^GFAP^+^ cells / total cells, n = 4 vs cKO = 0.42 ± 0.04 S100β^+^GFAP^+^ cells / total cells, n = 4; p < 0.05, unpaired T-Test) as well as GFAP^+^S100β^−^ B1 cells (Con = 0.033 ± 0.008 GFAP^+^S100β^−^ cells / total cells, n = 4 vs cKO = 0.096 ± 0.016 GFAP^+^S100β^−^ cells / total cells, n = 4; p < 0.05, unpaired T-Test) (Figure 2D). These data demonstrate a transition in the cellular composition at the ventricular surface. In the control animals there are rare cell bodies that express GFAP at the surface while, in the animals in which *Cyfip1* has been deleted, there is a larger proportion of cells expressing GFAP, with or without expression of S100β. This is indicative of an increase in the number of B1 cells at the surface and suggests either a translocation of cells from beneath the ventricular surface to the ventricular surface, or an expansion of the population of cells at the surface, or both. There is also an increase in the number of S100β^+^ GFAP^+^ astrocytes at the surface.

### Loss of Cyfip1 increases proliferating cells at the ventricular surface of adult animals

We hypothesized that the increase in GFAP^+^ cells at the ventricular surface is the result of the GFAP expressing NSCs translocating to the ventricular surface and dividing there. To test this hypothesis, we performed 5-ethynyl-2’-deoxyuridine (EDU) incorporation experiments to label actively cycling cells in the S phase. We injected 56 to 70 day old control animals as well as littermate cKO animals with a single intraperitoneal dose of 200 mg/kg of EDU. After 24 hours, animals were perfused with 4%PFA and whole mount and coronal sections were prepared for immunostaining, EDU labeling and quantification. Figure 3A demonstrates example whole mount preparations from control and cKO animals. We found a significant increase in the number of EDU^+^ nuclei at the ventricular surface in the cKO animals compared to controls (Con = 316 ± 67 cells / mm^2^, n = 3 animals vs 782 ± 49 cells / mm^2^, n = 3 animals; p < 0.05; unpaired T-Test) (Figure 3B), indicating an increase in proliferation of cells at the ventricular surface.

As the whole mount preparations only allow us to visualize the lateral wall of the lateral ventricle, we also examined coronal preparations of 56 to 70 day-old animals 24 hours after EDU injection. It is notable that the ventricles appear to be larger in the cKO animals and more cKO animals have an open configuration compared to the controls (Figure 3C). However, when we quantified the perimeter of the ventricles there was only a trend towards an increase in the perimeter of the cKO ventricle sections (Con = 3198 ± 184 µm / section; n = 4 vs cK0 = 3746 ± 340 µm/section; n = 3; p = 0.1868, unpaired T-Test; Data not shown).

We quantified the number of EDU^+^ cells at the ventricular surface by counting the total number of EDU^+^ cells lining the lateral wall, the medial wall, and the dorsal wall of the ventricle using every 6^th^ coronal section from the first section containing the anterior SVZ to the mid to posterior SVZ at the level of the dentate gyrus, as well as in the rostral migratory stream (RMS) just anterior to the ventricles (Figure 3C and 3D). As was found in the whole mount preparations, there was an increase in EDU^+^ cells in the cKO animals compared to the controls (Con = 257 ± 45 cells per hemisphere; n = 4 animals vs cKO = 714 ± 128 cells per hemisphere; n = 3 animals, p < 0.05; unpaired T-Test) (Figure 3D). This increase was reflective of a significant increase in proliferating cells in the lateral wall, the dorsal wall, and a trend towards an increase in proliferating cells in the medial wall (Figure 3D). When the rostral migratory stream was independently quantified there was also an increase in the number of EDU^+^ cells in the cKO vs the control (Figure 3C and 3F) (Con = 18 ± 13 cells per high powered field; n = 4 vs cKO = 124 ± 35 cells per high powered field, n = 3; p < 0.005, unpaired T-Test). This increase in the number of EDU^+^ cells entering the proximal RMS supports the hypothesis that there is an increase in the generation of new cells rather than only an increase in the number of cells that are being retained or failing to migrate from the ventricular surface. Furthermore, when we quantified the number of EDU^+^ cells that expressed GFAP on the cell surface, the relative proportion of GFAP^+^ EDU^+^ among all EDU^+^ cells is similar between the control and the cKO (Con = 0.89 ± 0.03 GFAP^+^EDU^+^ cells / total EDU^+^ cells; n = 4 vs cKO = 0.89 ± 0.05 GFAP^+^EDU^+^ cells / total EDU^+^ cells; p = 0.97; unpaired T-Test). This supports the hypothesis that it is a specific expansion of the GFAP-expressing B1 cell population rather than the addition of another proliferating population at the ventricular surface that results in this phenomenon.

### Acute loss of Cyfip1 in the adult SVZ disrupts niche architecture and alters NSC proliferation

Studies up to this point have examined the effect of loss of Cyfip1 during embryonic development and therefore cannot distinguish between the downstream effects of altering the embryonic neurogenic niche or a persistent need for Cyfip1 in the adult niche. To determine whether Cyfip1 plays a persistent functional role in the adult neurogenic niche, we developed an inducible conditional knockout animal (icKO) to delete the *Cyfip1* gene specifically in the NSCs in the subventricular zone of adult animals after the niche is already established. We used a tamoxifen inducible C*re-lox* system in which expression of a Cre recombinase protein with an estrogen receptor motif (Cre-ER), is driven by the *Nestin* promotor (Balordi and Fishell, 2007) (Figure 4A). This model allows for both temporal and spatial control over gene deletion. To verify Cre expression and to label cells in which recombination occurred, *Nestin-creER* animals were crossed with the *mTmG* reporter mouse (Muzumdar et al., 2007).

**Figure 4.**
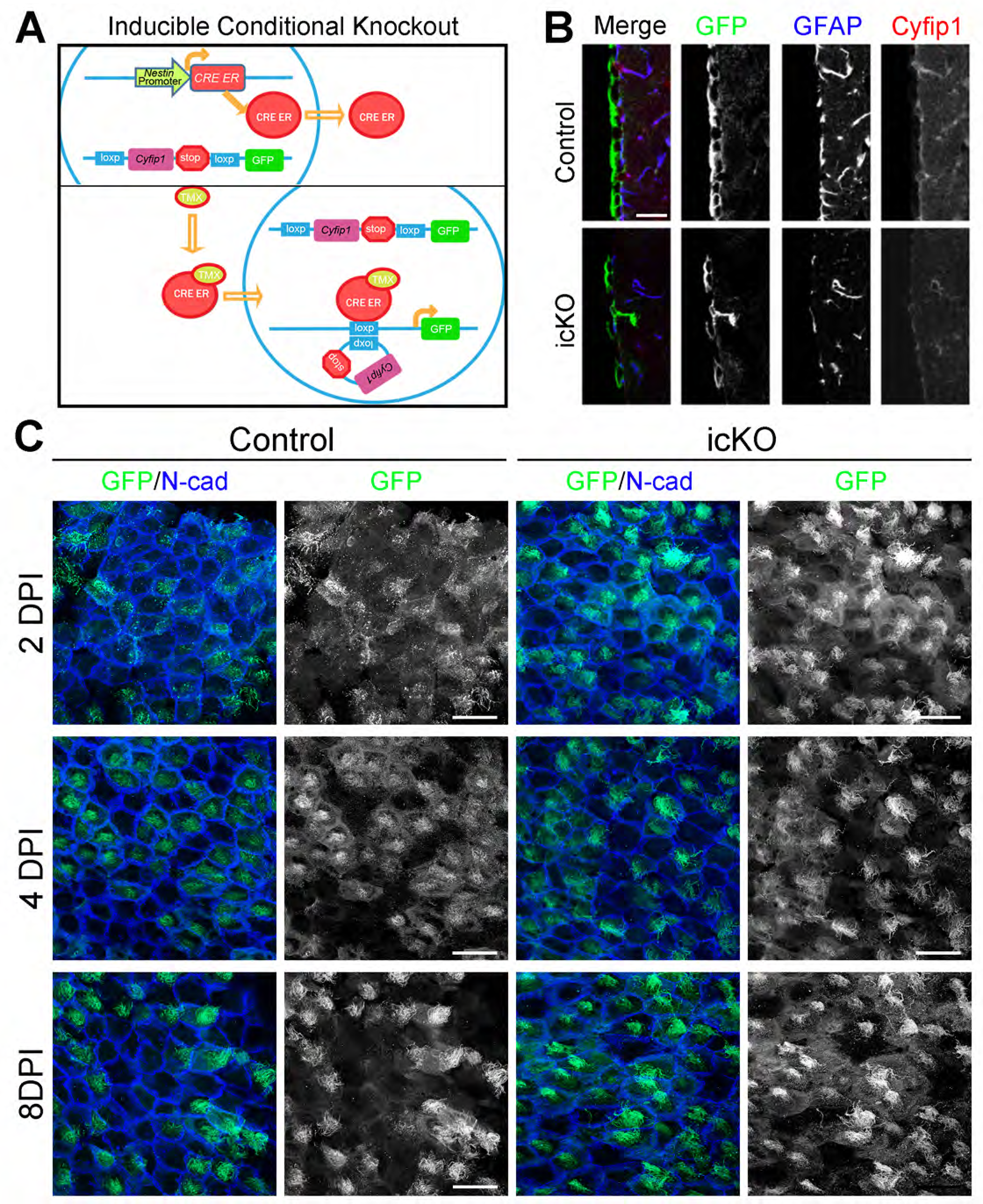
Acute deletion of *Cyfip1* in the adult SVZ. (A) Schematic illustration of *the Nestin-CreER:Cyfip1*^*f/f*^:*mtmg* inducible conditional knockout (icKO) animals. Tamoxifen injection results in nuclear translocation of the Cre-ER recombinase molecule resulting in deletion of *Cyfip1*. (B) Example ventricular wall of control and icKO animals demonstrating decreased Cyfip1 levels by immunofluorescence. Scale bar, 20 µm. (C) N-cadherin (blue) and GFP (green) immunofluorescence at 2, 4 and 8 days post injection (DPI) in whole mount preparations in control and icKO mice. Images taken with a 63X objective. Scale bars, 20 µm.

Adult control animals containing the *Nestin-cre*ER:*mTmG* transgenes that were wild type for *Cyfip1*, as well as icKO, animals with a *Nestin-cre*ER:*mTmG:Cyfip1*^f/f^ genotype were injected with tamoxifen between P56 and P84. Animals were then sacrificed at 2, 4 and 8 days post injection for analysis. Animals sacrificed at 8 DPI demonstrated decreased levels of Cyfip1 in the GFAP^+^ cells beneath the cell surface (Figure 4B). When the relative immunofluorescence for Cyfip1 staining was quantified, it was approximately 45% of controls (mean Cyfip1 immunofluorescence intensity 49.5 ± 5.5 intensity units, n = 5 cells from 1 Con animal vs 22.2 ± 4.1, n = 13 cells from 2 icKO animals; p = 0.002; unpaired T-Test). Whole mount staining with antibodies targeted against GFP and N-cadherin indicates that as early as 2 days post injection (DPI) Cre mediated recombination occurs in ependymal and B1 cells based on morphology at similar frequencies in both the control and icKO animals. There is an increase in the intensity of GFP immunofluorescence by 4 DPI and GFP^+^ cells remain at the surface at 8 DPI in both conditions (Figure 4C).

To determine whether Cyfip1 is required for regulation of the SVZ niche, we examined GFAP expression as well as N-cadherin expression at the ventricular surface at 2, 4 and 8 DPI. At 2 DPI, GFAP and N-cadherin staining is similar in the control and icKO. At 8 DPI, there is a marked increase in the number of GFAP^+^ cells at the cell surface (Figure 5A). This increase in GFAP immunoreactivity occurs in the form of an increased number of apical process clusters as well as an increased number of cell bodies at the cell surface and is reflective of a statistically significant increase in the number of normal and abnormal pinwheel formations (Mean of differences = 7.95 ± 1.37 GFAP^+^ cells / 100 mm^2^, p < 0.05, two tailed paired T test). Additionally, in the icKO whole mounts, while the GFAP^+^ processes project to the surface, they lack the tight pinwheel hub seen in control animals (Figure 5C arrowheads). When the expression of N-cadherin and GFAP in the pinwheel formations was examined at a high magnification, we saw that in the control SVZ there was a distinct demarcation between the GFAP-expressing apical projections and the ependymal cells at the surface and there was very little overlap between GFAP and N-cadherin immunofluorescence (Figure 5C, Control, arrowheads). However, there was intense N-cadherin staining surrounding the central apical projections. This is in stark contrast to the inducible knockout animals at 8 DPI, where there is a marked overlap in N-cadherin and GFAP expression. In the absence of Cyfip1, N-cadherin is no longer excluded from the center of the apical projection and there is no longer a clear demarcation between the B1 cells and the non-GFAP expressing cells on the surface (Figure 5C, 8 DPI, arrowheads). Additionally, the cell-cell junctions of the GFAP^−^ cells along the surface are thicker and less clearly defined compared to the control cells. At 2 and 5 DPI, some of the GFAP projections of the icKO animals are similar to the control animals. In others, the phenotype is similar to the GFAP^+^ processes of the 8 DPI icKO animals, suggesting that the process begins prior to the 8 DPI time point.

**Figure 5.**
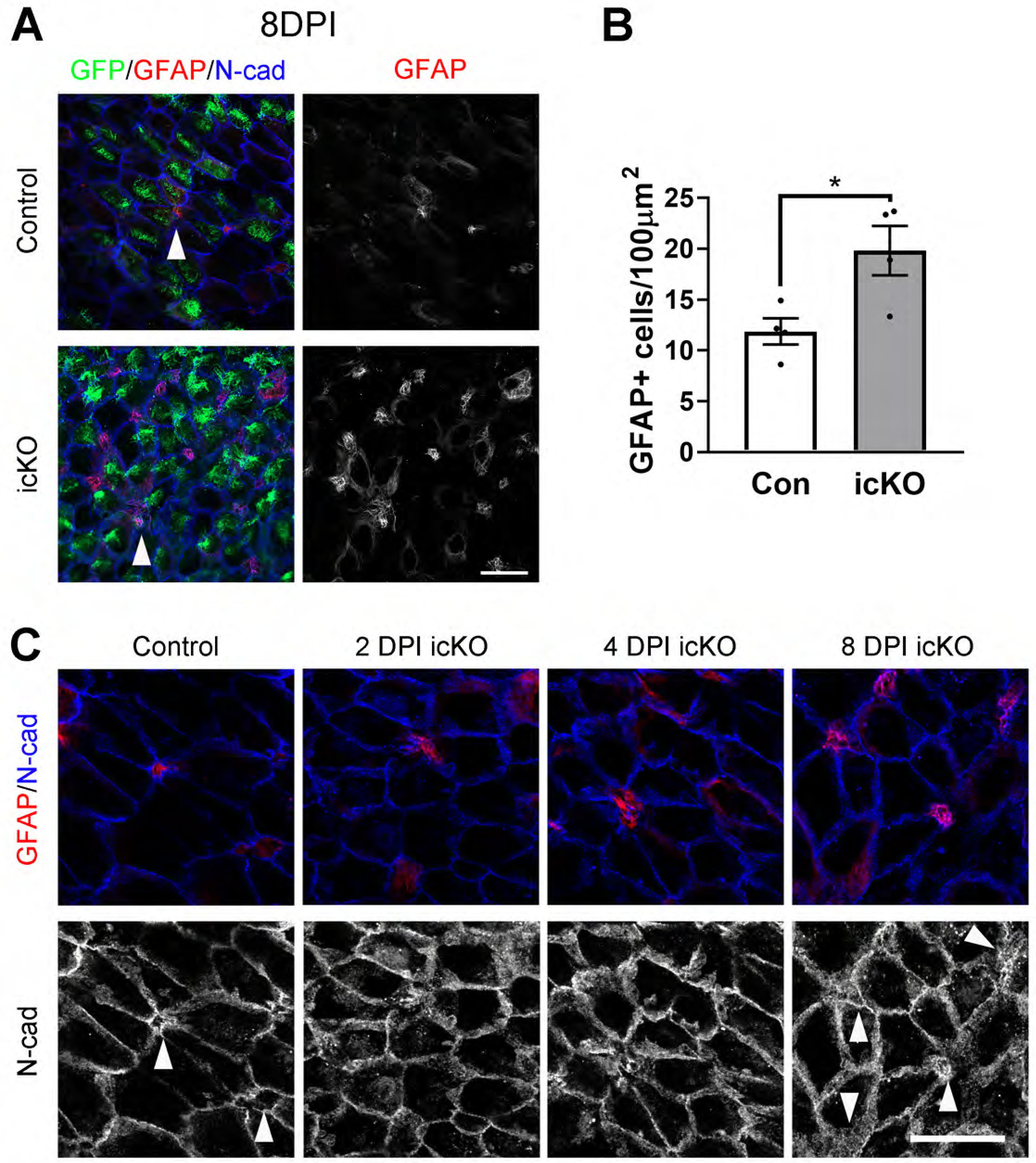
Acute loss of Cyfip1 disrupts the adult ventricular surface. (A) Whole mount preparations from control and induced conditional knockout (icKO) animals at 8 days post injection (DPI) immunostained for GFP (green), GFAP (red) and N-Cadherin (blue). Scale bar, 20 µm. (B) Quantification of the number of GFAP^+^ contacts with the cell surface of whole mount preparations. Mean of differences = 7.95 ± 1.37 GFAP^+^ cells / 100 mm^2^, n = 4 animals per condition. p < 0.05; two-tailed paired t-test. (C) High magnification images of whole mount preparations stained with GFAP (red) and N-Cadherin (blue). Arrowheads indicate type B1-cells and ependymal junctions in control versus cKO animals at 2, 4, and 8 days post injection. Scale bar, 20 µm. All images taken with a 63X objective.

We further examined the effect of acute Cyfip1 deletion on the localization of type B1 cells in coronal sections. We immunolabeled coronal sections from 8 DPI animals with antibodies directed against GFP (recombined mTmG^+^ cells) as well as anti-GFAP and S100β antibodies and determined the relative number of cells at the ventricular surface that expressed either or both proteins (Figure 6A and 6B). Similar to what was seen in the conditional knockout model (cKO) in which *Cyfip1* is deleted from the embryonic NSCs, we found that there is a significant increase in the number of GFAP^+^ S100β^−^ cells relative to total cells lining the surface (Figure 6B) (Con = 0.142 ± 0.031 GFAP^+^S100β^−^ cells / total cells, n = 4 vs icKO = 0.283 ± 0.042 GFAP^+^S100β^−^ cells / total cells, n = 3; p < 0.05; unpaired T-Test). This increased GFAP^+^ S100β^−^ population suggests that there is an increase in the number of type B1 cells at the ventricular surface. To confirm this, we immunostained for Sox-2, a transcription factor expressed in B1 cells and found a proportional increase in the number of Sox-2^+^ GFP^+^ cells relative to all GFP^+^ cells in the icKO animals compared to the controls (Con = 0.59 ± 0.02 Sox2^+^GFP^+^/ GFP^+^ cells, n = 3 vs icKO = 0.79 ± 0.04 Sox2^+^GFP^+^/ GFP^+^ cells, n = 3; p = 0.01; two tailed unpaired T-test) (Figure 6C and 6D). This indicates that of the recombined GFP^+^ cells, a higher proportion of cells either remain or become Sox2^+^ B1 cells. Additionally, Sox2^+^ cells were more prominent at the ventricular surface in the icKO brains compared to their subventricular localization in the control brains (Figure 6C). Together with the increase in the number of GFAP^+^ cells at the cell surface, this suggests either an increase in the proliferation of B1 cells or an increase in the translocation of the B1 cells to the surface. To distinguish between these possibilities, we examined the number of proliferating Sox2^+^ cells based on acute EDU-labeled cells (2 Hrs. post injection) that were also Sox2^+^GFP^+^ (Figure 6C and 6E) and compared them to the total number of GFP^+^ cells. We found a significant increase in the number of EDU-labeled Sox2^+^GFP^+^ cells in the icKO animals compared to controls (Figure 6E) (Con = 0.20 ± 0.02 EDU^+^Sox2^+^GFP^+^/ GFP^+^ cells, n = 3 vs icKO = 0.31 ± 0.03 EDU^+^Sox2^+^GFP^+^/ GFP^+^ cells, n = 3; p = 0.02; two tailed unpaired T-test). Taken together, these data indicate a specific ventricular translocation and expansion of the B1 NSCs as a result of acute loss of Cyfip1 in the adult SVZ.

**Figure 6.**
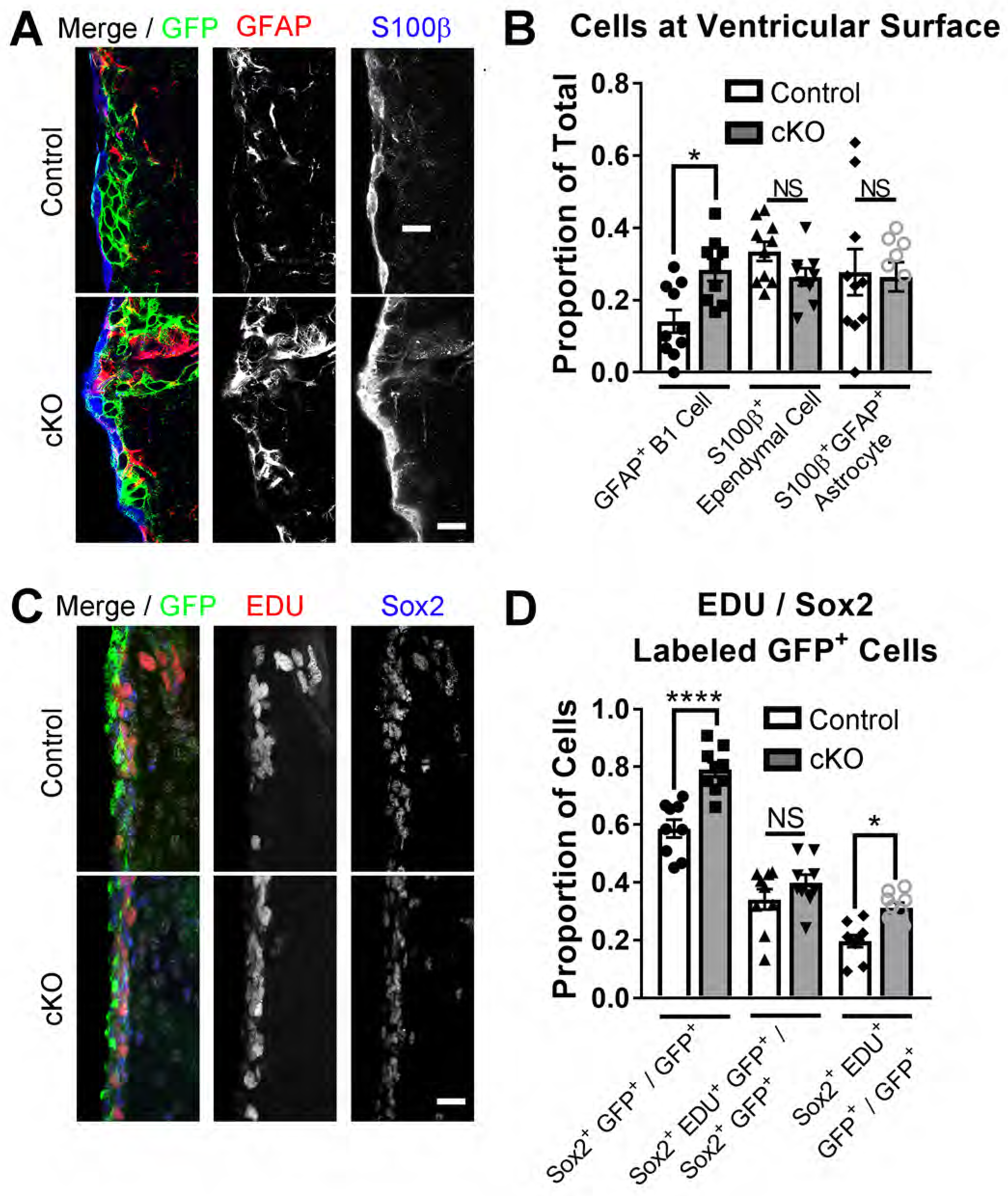
Acute loss of Cyfip1 increases proliferation in the ventricular and subventricular zone. (A) Example coronal sections immunostained for GFAP (red), MTMG (green – GFP), S100β (blue) from control and induced conditional knockout (icKO) animals. Scale bars, 10 µm. Images obtained with a 63X objective. (B) Quantification of GFAP^+^ and S100β^+^ cells compared to total MTMG^+^ (GFP^+^) cells at the ventricular surface of the lateral wall of the lateral ventricle. Values represent mean ± s.e.m. (n = 10 control and 9 icKO sections from n = 3 animals each condition; *p < 0.05; NS: p > 0.05. One way ANOVA, Sidak’s multiple comparisons test). (C) Example immunofluorescent images from control and icKO coronal sections stained against EDU (red), Sox2 (blue), GFP (green) and DAPI (grey – merged figure). Scale bar, 20 µm. Images obtained with a 40X objective. (D) Quantification of Sox2^+^ GFP^+^ and of Sox2^+^ EDU^+^ GFP^+^ cells compared to total number of GFP^+^ cells in control and knockout animals. Values represent mean ± s.e.m. (n = 9 sections each from n = 3 control and 3 icKO animals; *p < 0.0001; NS: p > 0.05; One way ANOVA, Sidak’s multiple comparisons test).

## DISCUSSION

In this study, we demonstrate that Cyfip1 is important for the proper establishment and maintenance of the adult SVZ niche architecture and the regulation of type B1 cell proliferation and localization. While the importance of Cyfip1 in embryonic development as well as mature neuronal plasticity is beginning to be appreciated (Abekhoukh and Bardoni, 2014; Abekhoukh et al., 2017; Yoon et al., 2014; De Rubeis et al., 2013a), this study is the first to suggest that Cyfip1 is a critical component for establishing and maintaining the adult NSC niche and regulating NSC fate. Our study further suggests that type B1 adult NSCs maintain the capacity for symmetric self-renewal to amplify their pool in the adult brain.

In contrast to the embryonic period, where there is prominent Cyfip1 expression in the apical membranes of the radial glial cells covering the entire ventricular surface (Yoon et al., 2014), our study demonstrates that the overall expression of Cyfip1 at the ventricular surface decreases in the adult SVZ as radial glial cells (RGCs) differentiate to ependymal cells. Remarkably, this indicates a specificity for NSCs as Cyfip1 continues to be expressed in the GFAP-expressing type B1 NSCs and is not prominent in the mature ependymal cells expressing S100β without GFAP. Similar to what is seen in the RGCs of embryonic development, there is specific localization of this protein to the apical processes at the ventricular surface in the adult SVZ and overlap with N-cadherin expression at cell-cell junctions. This indicates that, similar to embryonic development, Cyfip1 is involved in the regulation of adherens junctions in the adult SVZ and supports the hypothesis that Cyfip1 is required for niche maintenance.

Interestingly, Mirzedah et al (2008) have previously shown by electron microscopy that the adherens junctions in the pinwheel formations of the adult SVZ are asymmetric between ependymal cells and type B1 cells. Junctions between B1 cells are similar to those seen between RGCs in development. Ependymal – ependymal cell junctions are different from both (Mirzadeh et al., 2008). Asymmetric persistence of Cyfip1 expression and resultant differential regulation of adherens junctions in the B1 cells but not in ependymal cells is one way that this could be brought about. In support of this hypothesis, in the control SVZ, there is a discrete localization of N-cadherin to the cell-cell junctions in B1 cells. Acute deletion of *Cyfip1*, results in a dispersion of N–cadherin such that a discrete apical membrane ring surrounding the GFAP^+^ processes is not present. This suggests that Cyfip1 stabilizes N-cadherin at the apical cell-cell junction.

In the embryonic NSC niche, disrupted adherens junction stability leads to shorter cell cycles and a reduction of cells that exit the cell cycle (Gil-Sanz et al., 2014). Here, we see an increase in cell division as well as an increase in the number of B1 cells in the adult SVZ as a consequence of loss of Cyfip1 in the embryonic SVZ. Previous work has demonstrated that B1 cell divisions in the adult SVZ are either symmetrically self-depleting or symmetrically self-renewing and, over time, the balance between the two favors depletion. This leads to a progressive decrease in B1 cells with aging (Obernier et al., 2018). An increase in cell divisions can either lead to depletion or expansion of the overall NSC pool depending on which type of division is enhanced. In another model examining niche regulation of B1 cell division in the adult SVZ, loss of apical end feet anchoring in the niche by blocking vascular molecular adhesion molecule -1 (VCAM-1), leads to disrupted pinwheel architecture and increased self-depleting neurogenic divisions with a resultant depletion of the B1 cells (Kokovay et al., 2012). In contrast, in this study, we see increased proliferation and an expansion of B1 cells at the surface indicating an increase in symmetric self-renewing proliferation as a result of loss of Cyfip1. This result suggests that it is possible to attenuate or reverse the progressive depletion of B1 cells that is seen in the adult SVZ of control animals.

It is possible that, because the adult SVZ niche architecture is established at the end of embryonic development, loss of *Cyfip1* during embryonic development alters the structure of the niche and it is the dysregulated niche and not a persistent need for Cyfip1 in the adult niche that is important. However, the marked loss of localization of N-cadherin to cell–cell junctions, accompanied by the translocation of type B1 cells to the surface, and an increase in the proliferation of B1 cells when Cyfip1 is acutely depleted in our inducible conditional (icKO) model indicates that persistent Cyfip1 expression in B cells is indeed required to maintain the niche. Further, upregulation of self-renewing proliferation in the absence of Cyfip1 suggests that constitutive Cyfip1 expression in B1 cells influences cell fate decisions to favor symmetrically depleting divisions or maintenance of the quiescent state. The exact mechanisms by which Cyfip1 regulates these processes are unclear. It is possible that the symmetric vs. asymmetric adherens junctions provide information to the B1 cells about the surrounding cells and the loss of adhesion acts as a signal to the B1 cell to generate new cells through division. Alternatively, Cyfip1 may regulate cell fate choice through a signaling mechanism independent of its role in adherens junction maintenance and further studies are needed to elucidate which of these hypotheses is correct.

We demonstrate that B1 cells in the adult can reactivate their capacity for symmetric self-renewing divisions after embryonic development. This could have implications for regeneration as well as oncologic transformation and with regards to the latter possibility, it should be noted that CYFIP1 has been proposed as a tumor invasion suppressor in humans (Silva, J. M. et al., 2009). Additionally, the phenotype observed in our cKO model demonstrating increased symmetric renewing divisions in the adult after embryonic deletion is pertinent to recent findings demonstrating that humans who are haploinsufficient for *CYFIP1* due to deletion of the 15q11.2 locus, where the gene is located, demonstrate microstructural alterations in white matter on MRI (Silva, A. I. et al., 2019) and that mice that are haploinsufficient for *Cyfip1* have decreased myelination in of the corpus callosum, decreased numbers of oligodendrocytes and abnormal behavior (Silva, A. I. et al., 2019; Dominguez-Iturza et al., 2019). Although there are many hypotheses as to why loss of Cyfip1 in mice could alter myelination based on its known role in actin nucleation which is needed for migration and adhesion, the data presented here suggest that the increased symmetric B1 cell renewing divisions occur at the expense of the generation of oligodendrocytes, resulting in impaired myelination either in the pre- or postnatal period or both. Finally, in the cKO animal, but not the acute icKO model, an additional consequence of the loss of *Cyfip1* and increased B1 cell generation is that there are increased double GFAP^+^ S100β^+^ cells at the ventricular surface that likely represent mature astrocytes. This indicates that one consequence of this shift in the mode of cell divisions is an eventual increase in astrogenesis which may have additional consequences.

CYFIP1 is located in the 15q11.2 locus in humans and deletions or duplications in this region are found in patients with epilepsy, intellectual disability (ID), autism and schizophrenia, suggesting that CYFIP1 is important for neural development (van der Zwaag et al., 2010; von der Lippe et al., 2011; Doornbos et al., 2009; Borlot et al., 2017; Mullen et al., 2013; Mefford et al., 2010; de Kovel et al., 2010; Rudd et al., 2014). Copy number variation in the 15q11.2 locus also results in changes in white matter microstructure (Silva, A. I. et al., 2019). The role of Cyfip1 as a member of the WAVE regulatory complex (WRC) in regulating actin nucleation makes it an ideal candidate to regulate synaptic plasticity as well as a regulator of early neural development (De Rubeis et al., 2013b; Abekhoukh et al., 2017; Yoon et al., 2014). Results presented here suggest that it continues to be important in postnatal NSC regulation with important downstream effects on postnatal neuron and oligodendrocyte genesis. This study builds on a previous study (Yoon et al., 2014) demonstrating the necessity of Cyfip1 for the establishment of apical basal polarity in embryonic neurogenesis and establishes a persistent requirement for its expression in the adult neurogenic niche. These studies indicate that Cyfip1 is crucial to NSC behavior and the neurogenic niche throughout life. Importantly, we show here that Cyfip1 is required to suppress self-renewing B1 cell divisions and that the adult NSC pool can be reactivated to favor self-renewal, even in the adult SVZ.

## Acknowledgements

This work was supported by grants from NIH (R25NS065729 and K12NS098482 to C.W.H., R37NS047344 to H.S., R35NS097370 to G-l.M) and DOD (W81XWH1810174 ALSRP to N.J.M.). The *Nestin-CreER* animals were kindly provided by Gordon Fishell. We are grateful to the lab members of the Song and Ming laboratories for feedback and technical support, in particular Kim Christian, Daniel Berg, Allison Bond, Tong Ma, Yi Zhou, Fadi Jacob and Jordan Schnoll.

